## Supplementary Material and Methods for "A gradient of Wnt activity positions the neurosensory domains of the inner ear"

**Plasmids**

Plasmids used and generated for the study are listed in the table below.

| Plasmid name | Plasmid type | Insert | Promoter | References |
| --- | --- | --- | --- | --- |
| 5TCF::H2B-RFP | Wnt reporter  (TopRed) | 5 x TCF/Lef binding sites; H2B-RFP fusion protein | Minimal TK | 5xTCF-bs-RFP ^1^ |
| T2-5TCF::nd2Scarlet | Wnt reporter  (Tol2) | 5 x TCF/Lef binding sites; nuclear-localized and destabilized Scarlet | Minimal TK | This study and ^2^ |
| T2-Hes5::nd2EGFP | Notch reporter  (Tol2) | mouse Hes5 promoter; nuclear-localized and destabilized EGFP | Mouse Hes5 | ^3^ |
| Hes5::d2FP635 | Notch reporter  (Slax) | mouse Hes5 promoter; destabilized  Turbo FP635 | Mouse Hes5 | ^3^ |
| RCAS-βcat-LOF | β-catenin LOF  (RCAS) | HA-tagged truncated form of Xenopus  β-catenin (lacking 134aa in C-terminus and 147aa at the N-terminus) | LTR | RCAS/*β-catenin ^4^  Truncated Xenopus  β-catenin « T5 » construct in ^5^ |
| T2-βcat-LOF | β-catenin LOF  (Tol2) | Membrane-localized Cherry; 2A self-cleaving peptide; triple HA-tagged truncated form of Xenopus β-catenin (subcloned from RCAS-βcat-LOF) | CAG | This study |
| PB-βcat-GOF | β-catenin GOF  (PiggyBac) | S33Y* mutated form of human β-catenin; IRES; H2B-EGFP fusion protein | CAG | PiggyBac-CAG-β-catenin^S33Y^-IRES-EGFP ^1^ |
| T2-βcat-GOF | β-catenin GOF  (Tol2) | S33Y* mutated form of human β-catenin; IRES; H2B-EGFP fusion protein | CAG | This study |
| pNICD1-EGFP | Notch GOF  (pCAGGS) | HA-tagged chicken Notch1 intracellular domain; IRES; EGFP | CAG | NICD1-IRES-GFP ^6^ |
| pDN-MAML1-EGFP | Notch LOF  (pCAGGS) | Truncated, dominant-negative form of human Mastermind-like 1 fused to EGFP | CAG | CAGGS-DN-MAML1–EGFP ^7^ |
| T2-EGFP | Control  (Tol2) | Enhanced Green Fluorescent Protein | CAG | pT2K-CAGGS-EGFP ^8^ |
| T2-mEGFP | Control  (Tol2) | Membrane-localized Enhanced Green Fluorescent Protein | CAG | This study |
| T2-mRFP | Control  (Tol2) | Membrane-localized Cherry | CAG | This study |
| mPB | PiggyBac transposase  (pCAGGS) | PiggyBac transposase | CAG | ^9^ |
| pCAGGS-T2-TP | Tol2 transposase  (pCAGGS) | Tol2 Transposase | CAG | ^8^ |

**Quantification of the Wnt gradient profile**

A schematic representing the pipeline for analysis of fluorescence intensity profile of Wnt reporter along the dorso-ventral axis of otocyst transfected with 5TCF::H2B-RFP.

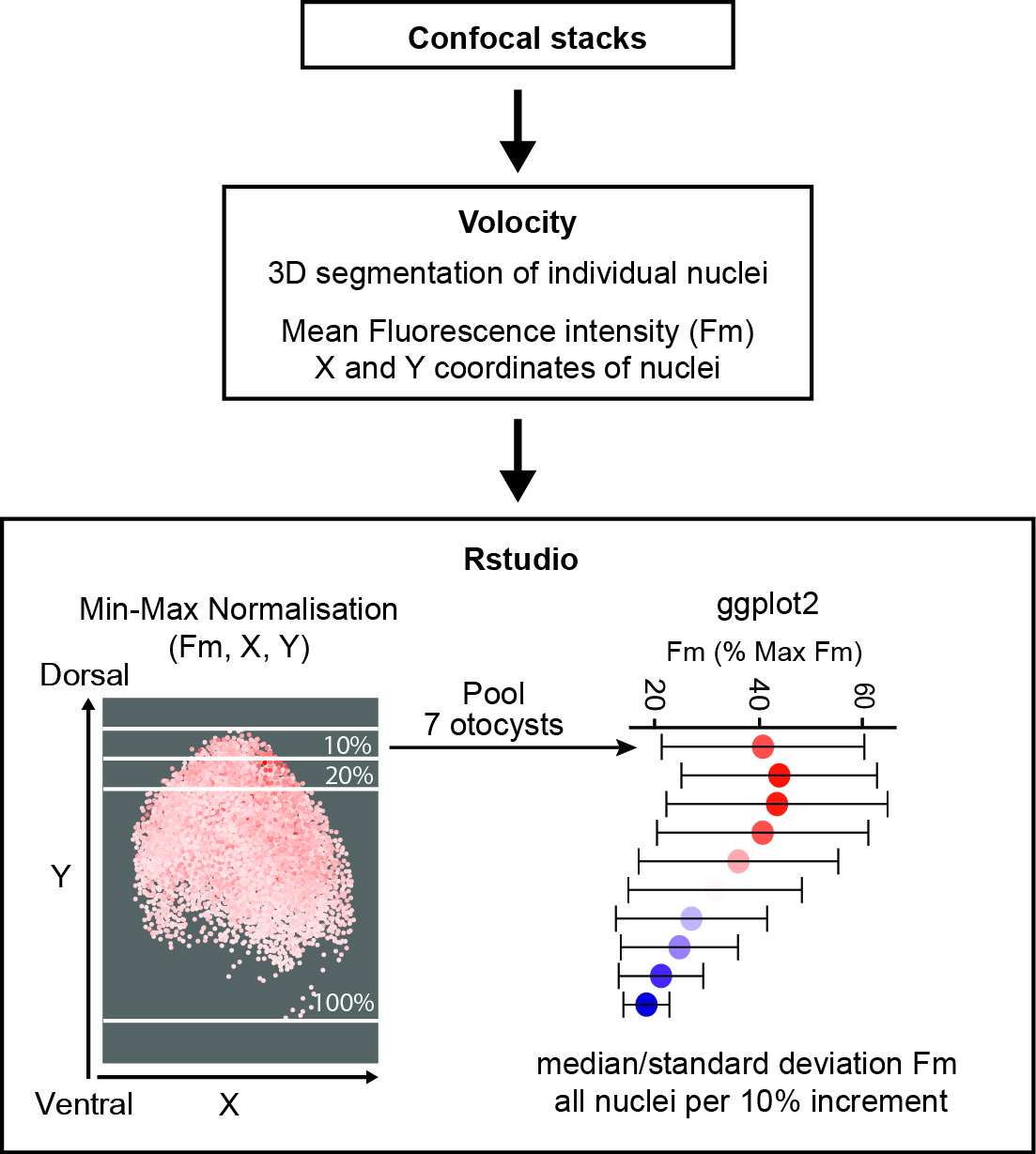

**Immunohistochemistry**

Sample dissection protocol depended on the age of embryos. For E7 embryos, the heads were halved along the midline, the inner ear was dissected and the otic cartilage was removed from the basilar papilla and trimmed at the dorsal side to expose the otic epithelium. For embryos aged E3-E4, the embryo was dissected along the midline, the hindbrain was removed, and the region surrounding the otocyst was only partially trimmed to facilitate orientation. A small opening was made at the dorsal tip of the otocyst using a fine needle and the tissue was permeabilized in PBS containing 0.3% Triton and 10% goat serum for 30 min at room temperature. Specimens were incubated with primary antibodies diluted in 0.1% Triton in PBS at 4°C overnight. On the next day, tissues were rinsed with PBS at room temperature and incubated with secondary antibodies diluted in 0.1% Triton and 10% goat serum at 4°C overnight. Afterward, tissues were again rinsed with PBS and mounted in Vectashield Antifade Mounting Medium (Vector laboratories). A fine layer of vacuum grease was applied between the slide and coverslip to avoid excessive flattening of the tissue.

**Determination of IWR-1 working concentration**

qPCR was used to establish a working concentration of IWR-1 in organotypic culture. E3 Otic explants (4-5 chicken otocysts per condition) were incubated in media containing 50µM, 150µM, and 300µM of IWR-1 or DMSO (vehicle) as a control. After 24h incubation, total RNA was isolated using the RNAqueous™-Micro Total RNA Isolation Kit (Ambion) and reverse transcribed using iScript cDNA Synthesis Kit (Bio-Rad). qPCR reactions were performed with Quantifast Syber Green (Qiagen). The effects of the treatment were analysed by testing the decrease in the expression levels of Lgr5 and Axin2, two genes positively regulated by Wnt signalling ^10, 11^. The relative quantification of expression was analyzed using the ΔΔCt method ^12^ and showed the significant downregulation, 55% for Lgr5 and 78% for Axin2, at 300µM of IWR-1.

**RNA-Seq and bioinformatics analyses**

Total RNA was extracted from each individual otocysts using the RNAqueous™-Micro Total RNA Isolation Kit (Ambion) according to the manufacturers protocol. The quality of isolated RNA was tested using Agilent 2200 Tapestation and only samples with a value of RNA integrity number (RIN) of at least 9 were used for library preparation using SMART-Seq v4 Ultra Low Input RNA Kit (Clontech Laboratories, Inc.) by UCL Genomics. Briefly, cDNA libraries were generated using the SMART (Switching Mechanism at 5' End of RNA Template) technology using 10 cycles of PCR. cDNA was checked for integrity and quantity on the Agilent Bioanalyser using the High Sensitivity DNA kit, and 200pg of cDNA was then converted to sequencing library using the Nextera XT DNA protocol (Illumina, San Diego, US), which allows for tagmentation and sample indexing for multiplex sequencing. Samples were sequenced 43bp paired-end read and ~16M reads per sample length on NextSeq 500 instrument (Illumina, San Diego, CA, US). Run data were demultiplexed and converted into fastq files using Illumina’s bcl2fastq Conversion Software v2.19. Kallisto package (Bray, Pimentel et al. 2016) was used to map reads to a chicken reference genome (Gallus_gallus-5.0) and to quantifying abundances of transcripts. Differentially expressed genes were identify using Sleuth package (Pimentel, Bray et al. 2017) by directly comparing left and right ear from the same embryo. Genes with p-value < 0.05 were considered as significant. Functional annotations were downloaded from ENSEMBL using biomaRt and signalling pathway enrichment and Biological Function enrichment analysis were performed using Toppgene online tool with default settings.
